# Neuronal AMPK coordinates mitochondrial energy sensing and hypoxia resistance

**DOI:** 10.1101/2020.05.01.073007

**Authors:** Brandon J. Berry, Aksana Baldzizhar, Andrew P. Wojtovich

## Abstract

Organisms adapt to their environment through coordinated changes in mitochondrial function and metabolism. The mitochondrial protonmotive force (PMF) is an electrochemical gradient that powers ATP synthesis and adjusts metabolism to energetic demands via cellular signaling. It is unknown how or where transient PMF changes are sensed and signaled due to lack of precise spatiotemporal control *in vivo.* We addressed this by expressing a light-activated proton pump in mitochondria to spatiotemporally “turn off” mitochondrial function through PMF dissipation in tissues with light. We applied our construct – mitochondria-OFF (mtOFF) – to understand how metabolic status impacts hypoxia resistance, a response that relies on mitochondrial function. mtOFF activation induced starvation-like behavior mediated by AMP-activated protein kinase (AMPK). We found prophylactic mtOFF activation increased survival following hypoxia, and that protection relied on neuronal AMPK. Our study links spatiotemporal control of mitochondrial PMF to cellular metabolic changes that mediate behavior and stress resistance.

## INTRODUCTION

Mitochondria are organelles that provide energy to cells. Matching energy supply with energy demand is coordinated through various process and is critical for cellular adaptation and survival under changing conditions. Therefore, an organism’s health depends not only on mitochondrial energy production, but on the ability to sense metabolic status and the ability of mitochondria to signal appropriately to use that energy (To et al., 2019, McElroy and Chandel, 2017). The mechanisms through which cellular energy-related signaling can occur likely depend on the electrochemical gradient across the mitochondrial inner membrane (IM).

Mitochondrial respiration results in a proton gradient across the IM known as the protonmotive force (PMF). Ultimately, respiratory complexes of the electron transport chain (ETC) convert chemical energy into electrical potential energy by pumping protons across the IM, creating this gradient. The PMF can then be consumed at ATP synthase, converting the PMF back to chemical energy in the form of ATP to catalyze reactions (Figure 1A). This process is called oxidative phosphorylation, as ETC activity consumes oxygen to maintain PMF that is then used to phosphorylate ADP to ATP. The PMF is also used for substrate (Meisner et al., 1972) and protein (Pfanner and Geissler, 2001) import to the mitochondrial matrix or IM, calcium signaling (Pfanner and Geissler, 2001), and quality control management through autophagy (Lim et al., 2019, Palikaras et al., 2018), mitochondrial biogenesis (Li et al., 2017), and proteostatic responses that involve mitochondria-nucleus communication (Rolland et al., 2019).

**Figure 1.**
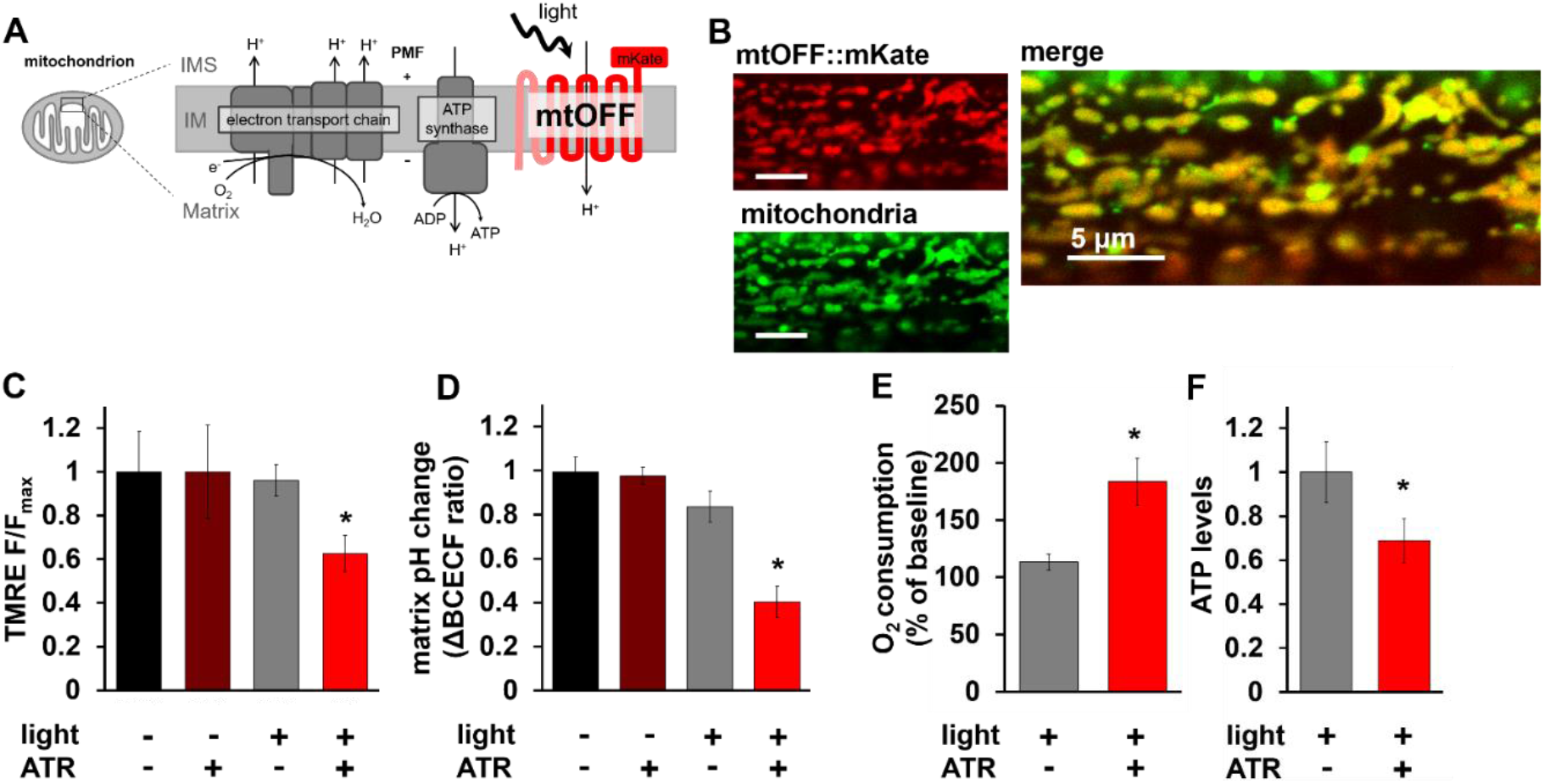
Mitochondria-OFF (mtOFF) decreases mitochondrial protonmotive force. **A)** Schematic of mtOFF targeted to the mitochondrial inner membrane (IM) to dissipate the protonmotive force (PMF). Electron transport chain (ETC) complexes together consume O_2_ and generate the PMF by proton (H^+^) pumping from the matrix to the intermembrane space (IMS). Mitochondrial ATP synthase uses the PMF to make ATP from ADP. The N terminal mitochondria targeting sequence and two transmembrane regions of the rat SDHC1 protein are shown in pink fused to the red proton pumping portion of mtOFF. The red fluorescent protein mKate is shown on the C terminus in the IMS in red. Light activation of mtOFF results in proton pumping from the IMS to the matrix. **B)** Fluorescent images show muscle mitochondria of a living *C. elegans* ubiquitously expressing mtOFF. Red signal shows mKate fluorescence and green signal shows MitoTracker Green staining of mitochondria. The merged image shows the mitochondrial localization of the mtOFF∷mKate construct overlapping with MitoTracker Green signal. Scale bars are 5 μm. **C)** Quantification of TMRE fluorescence intensity in isolated mitochondria incubated with succinate to fuel membrane potential (ΔΨ_m_) shows decreased ΔΨ_m_ upon mtOFF activation. Data are normalized to dark conditions and full light doses are presented in Supplementary Figure 2B. One-way ANOVA was performed with Tukey’s test for multiple comparisons, *p = 0.0247. Data are means ± SEM, n = 4 independent mitochondrial isolations. **D)** Quantification of change in BCECF-AM ratio in isolated mitochondria fueled with succinate shows decreased mitochondrial matrix pH after mtOFF activation. One-way ANOVA was performed with Tukey’s test for multiple comparisons, -ATR -light vs +ATR +light *p = 0.0002, +ATR -light vs +ATR +light *p = 0.0002, -ATR +light vs +ATR +light *p = 0.0036. Data are means ± SEM, n = 4-5 independent mitochondria isolations. **E)** O_2_ consumption rates of whole animals normalized to dark conditions were increased upon mtOFF activation. Raw O_2_ consumption rates are shown in Supplementary Figure 2D. Two-tailed unpaired t test was performed, *p = 0.0195. Data are means ± SEM, n = 5, where one n is one O_2_ consumption rate from ~1500 animals in a Clark type O_2_ electrode. **F)** Relative ATP levels normalized to dark conditions from whole animals was decreased upon mtOFF activation. Two-tailed unpaired t test was performed, *p = 0.0230. Data are means ± SEM, n = 5 independent assays from three plates each for each condition containing at least 100 animals.

PMF changes are physiologic and adaptive, and function to alter cellular functions regulated by mitochondria. For example, organisms that experience different oxygen tensions can change ETC activity and PMF (Tello et al., 2011, Weinberg et al., 2000), and the ability to adapt to these changes is linked to survival (Yin et al., 2013, Kim et al., 2006). Inability to respond to changes through the PMF can result in pathology. For example, during hypoxia, the PMF is lost (Kaufman and Crowder, 2015) and leads to damage or death if not restored (Pena et al., 2016). Finding conditions that allow resistance to hypoxia is important to better understand diseases where oxygen (O_2_) deprivation causes cellular damage. During ischemic stroke, for example, neurons are starved of the O_2_ that fuels the ETC, resulting in decreased PMF. The PMF is rapidly restored upon reperfusion, and this rapid reestablishment can drive oxidative stress from an overwhelmed ETC (Onukwufor et al., 2019, Chouchani et al., 2014). This oxidative damage is highly sensitive to the PMF, where a slight decrease at reperfusion can greatly decrease oxidative stress (Lambert and Brand, 2004, Komlodi et al., 2018). Artificially decreasing the PMF with drugs is protective in hypoxia models, but spatial and temporal nuance is missing due to the imprecise nature of pharmacology. Pharmacologic approaches are not tissue or cell-type selective. In addition, once pharmacologic agents act to decrease PMF, they cannot be reversed. Therefore, using drugs to decrease the PMF is dangerous due to narrow therapeutic windows and lack of tissue selectivity (Dufayet et al., 2020, Goedeke et al., 2019, Bleasdale et al., 2018). This difficulty in developing PMF therapy highlights the need for spatiotemporal control in experimental models to understand how decreasing the PMF provides stress resistance. How and where the metabolic state of an organism is sensed, and how that translates to physiologic changes remains unclear. With this study, we sought to understand how and where animals sense PMF dissipation. We also asked how and where energy sensing elicits protective signaling prior to hypoxia exposure in living animals. We achieved this spatiotemporal control through optogenetics.

We have recently shown that a decreased PMF prior to pathologic hypoxia is necessary and sufficient for protective signaling in *Caenorhabditis elegans* (Berry et al., 2020b). Using the mitochondria-ON (mtON) construct we determined the temporal effect by using the acute reversibility of light exposure. Our result was supported by evidence in cells using a similar optogenetic approach to spatiotemporally control the PMF, showing brief PMF loss preconditions cells to be resistant to later, severe PMF disruption (Ernst et al., 2019). Decreasing the PMF during hypoxia to relieve oxidative stress is thoroughly characterized (Otani, 2004, Ozcan et al., 2013, Hoerter et al., 2004, Sack, 2006, Korde et al., 2005); here we address how decreasing the PMF prophylactically signals protection against impending hypoxic insults *in vivo* through energy sensing. We hypothesized that decreasing the PMF with high spatial and temporal precision using optogenetics would show how and where energetic signals are transmitted from mitochondria throughout an organism to mount a stress-resistance response to hypoxia.

Responses to energy availability may provide a mechanism of hypoxia resistance through changes in the PMF, as fuel for the ETC impacts how O_2_ is used. Indeed, energy sensing through food availability plays an important and specific role in responses to O_2_ deprivation in model organisms. For example, the nematode *C. elegans* requires AMP-activated protein kinase (AMPK) signaling for healthy recovery from hypoxia (LaRue and Padilla, 2011). *C. elegans* AMPK responds to nutritional state and coordinates hypoxia resistance (LaRue and Padilla, 2011). These results translate to mammalian models, as AMPK protects against hypoxic pathologies like ischemic reperfusion injury (Wang et al., 2017, Kim et al., 2011). In addition, AMPK acts as a neuronal energy sensor capable of changing an animal’s behavior based on food availability (Ahmadi and Roy, 2016, Yurgel et al., 2019). These combined data lead us to ask how mitochondrial energy status is sensed and how it is translated to hypoxia resistance.

In our previous work with mtON, energy sensing signaling through AMPK was silenced when mitochondrial function was increased with light (Berry et al., 2020b). Here, we causally link AMPK signaling as a direct readout of changing PMF and confirm its requirement for resistance to hypoxia. Given that neuronal AMPK can control an organism’s behavior in response to sensed energy availability (Ahmadi and Roy, 2016), we show that neuronal AMPK is regulated downstream of changes in the PMF. Further, we show that neuronal AMPK activation downstream of dissipated PMF is sufficient for hypoxia resistance, linking perceived internal energy-state to stress-resistance through metabolism.

## RESULTS

### Mitochondria-OFF (mtOFF) is expressed in *C. elegans* mitochondria

Recently, light-activated channels called channelrhodopsin 2 (ChR2) have been directed to mitochondria to dissipate the protonmotive force (PMF) in response to light (Ernst et al., 2019, Tkatch et al., 2017). This technique allows light-specific dissipation of PMF and has been validated in different metabolic contexts to impact broad cellular signaling activities (Ernst et al., 2019, Tkatch et al., 2017). We extend this approach using the light-activated proton pump from mitochondria-ON (mtON) in our previous work (Berry et al., 2020b), here oriented to dissipate the PMF in whole animals. Using a light-activated proton pump required yellow-green light and avoided the use of blue light that is required for ChR2. Blue light can be damaging to biological samples (De Magalhaes Filho et al., 2018), especially mitochondria, which contain light-sensitive redox-active flavins that could alter oxidative stress (Yang et al., 2017, Trewin et al., 2018). Proton selectivity (Waschuk et al., 2005) also avoids caveats of ChR2 moving other cellular ions such as potassium and calcium (Schneider et al., 2013, Nagel et al., 2003), both ions that are important for other mitochondrial and cellular functions (Bartok et al., 2019, Paggio et al., 2019). Using a ubiquitously expressed gene promoter, we expressed the light-activated proton pump, Mac (Waschuk et al., 2005), in *C. elegans* mitochondria. Our construct was oriented to pump protons from the mitochondrial intermembrane space (IMS) into the matrix to dissipate the PMF (Figure 1A). We called this construct mitochondria-OFF, or mtOFF, due to its ability to “turn off” mitochondrial function through the PMF in response to light, as validated here and by other studies using ChR2 (Ernst et al., 2019, Tkatch et al., 2017). The mitochondrial targeting sequence (MTS) and part of the coding sequence of the of the SDHC1 protein were used to direct and orient mitochondrial expression of Mac. This SDHC1∷Mac construct is the functional unit mtOFF (Figure 1A). We then used C terminal fusion to the red fluorescent protein mKate for visualization in living animals. Using CRISPR/Cas9 genome editing we integrated a single copy of the mtOFF construct into the *C. elegans* genome under the control of a ubiquitous promoter (*eft-3p*) using the mmCRISPi technique (Philip et al., 2019) to avoid over-expression artifacts. Fluorescence indicated mitochondrial expression when observed by confocal microscopy in animals stained with MitoTracker Green (Figure 1B). Further, in single mitochondria of living animals, C terminal mKate fluorescence was distant from MitoTracker Green fluorescence in the mitochondrial matrix, but overlapped with IMS targeted GFP (Supplementary Figure 1 A-C). In line with our previous work (Berry et al., 2020a), these data suggest the orientation of the proton pump was as predicted (Figure 1A) indicated by the distance between red and green fluorescence in each case (Supplementary Figure 1D).

### mtOFF decreases mitochondrial PMF in response to light

Using mtON, we previously showed that a proton pump oriented oppositely from our approach here was able to generate a PMF independent of ETC function (Berry et al., 2020a). Therefore, a pump inserted into the membrane in the opposite orientation is predicted to decrease the PMF. Based on the expected topology (Figure 1A) and the localization of mKate (Supplementary Figure 1A-D), we asked if mtOFF could decrease the PMF. We tested this by measuring the PMF in response to mtOFF activation in two independent assays. Optogenetic proteins such as the Mac component of mtOFF require a cofactor, all-*trans* retinal (ATR), for photocurrents to occur (Chow et al., 2010, Berry et al., 2020b). *C. elegans* require ATR supplementation due to endogenous absence in the organism (Husson et al., 2012). Therefore, each of our experiments controlled for ATR supplementation, as well as light exposure to control for any confounding effects of light or ATR. Only light exposure and the presence of ATR allow mtOFF function (Husson et al., 2012, Berry et al., 2020b). Under these conditions, mtOFF decreased the PMF in response to light when observed through both components of the gradient, the membrane potential (ΔΨ_m_) and the pH gradient (ΔpH) (Supplementary Figure 2A). Isolated mitochondria loaded with the ΔΨ_m_ fluorescent indicator tetramethylrhodamine ethyl ester (TMRE) were incubated with succinate to fuel respiration and to maintain the PMF. Upon light exposure, ΔΨ_m_ decreased significantly (Figure 1C) and light-dose dependently (Supplementary Figure 2B). The ΔpH was assessed by observing BCECF fluorescence in isolated mitochondria. BCECF-AM is a ratiometric pH indicator that can be loaded into isolated mitochondria (Jung et al., 1989, Aldakkak et al., 2010) to determine changes in matrix pH. When provided succinate to maintain PMF, mtOFF activation resulted in significant, reversible matrix pH decrease (Figure 1D & Supplementary Figure 2C), indicating decreased ΔpH. These data independently demonstrate that mtOFF activation dissipated the PMF through both its components, ΔΨ_m_ and ΔpH.

Respiratory control is a phenomenon of increased ETC activity in response to dissipated PMF (Nicholls, 1977, Mitchell, 2011). The ETC increases activity (and resulting O_2_ consumption) in attempt to maintain PMF. We therefore measured whole-animal O_2_ consumption rates and ATP levels to assess the consequences of dissipated PMF. mtOFF activation resulted in around a 70% increase in respiration compared to control conditions (Figure 1E & Supplementary Figure 2D), similar to results obtained in a cell model with mitochondria targeted ChR2 (Tkatch et al., 2017). We then measured relative ATP levels to assess if mtOFF was indeed interrupting respiratory control *in vivo.* Whole animal ATP levels were decreased after mtOFF activation (Figure 1F), indicating that mtOFF-mediated PMF dissipation decreased cellular energetics. Together, these results indicated that mtOFF functioned to decrease the PMF to alter the internal metabolic state of organisms.

### mtOFF modulates energy sensing behavior through neuronal AMPK

To understand how PMF dissipation affects metabolism and physiology in living animals, we used *C. elegans* expressing mtOFF. As we have previously shown, AMP-activated protein kinase (AMPK) activity is altered downstream of PMF changes and in signaling that affects *C. elegans* behavior (Berry et al., 2020b). *C. elegans* respond to food availability and their internal metabolic state (fed versus starved) (Fagan et al., 2018, Ryan et al., 2014) through AMPK by increasing or decreasing their locomotion speed (Sawin et al., 2000, Tsalik and Hobert, 2003). In the presence of food, animals will move slowly to stay in its presence, and in the absence of food, animals will increase their movement speed (Figure 2A&B). This behavior is blunted in animals lacking AMPK activity (*aak-2* mutant animals (Lee et al., 2008, Ahmadi and Roy, 2016, Berry et al., 2020b)) (Figure 2D). Previously, we found that increasing the PMF silenced AMPK signaling under starvation conditions, and could slow animal locomotion (Berry et al., 2020b). Conversely, mtOFF activation caused increased AMPK phosphorylation and therefore activation (Figure 2C&D), causing increased locomotion under fed conditions (Figure 2E&F). mtOFF was able to create an energy demand resulting in increased locomotion when animals were still in the presence of food. This optogenetic effect was abolished when AMPK was inhibited with compound C. The mtOFF-mediated increase in locomotion was also blocked in AMPK mutant animals expressing mtOFF (Figure 2F). No optogenetic effects were observed in conditions off of food, as a high energy demand is already present under starvation conditions (Supplementary Figure 3). These data suggest AMPK activity is regulated downstream of mitochondrial function to regulate metabolic demand, and they show that mtOFF can control animal behavior through the PMF and resultant energy sensing signaling.

**Figure 2.**
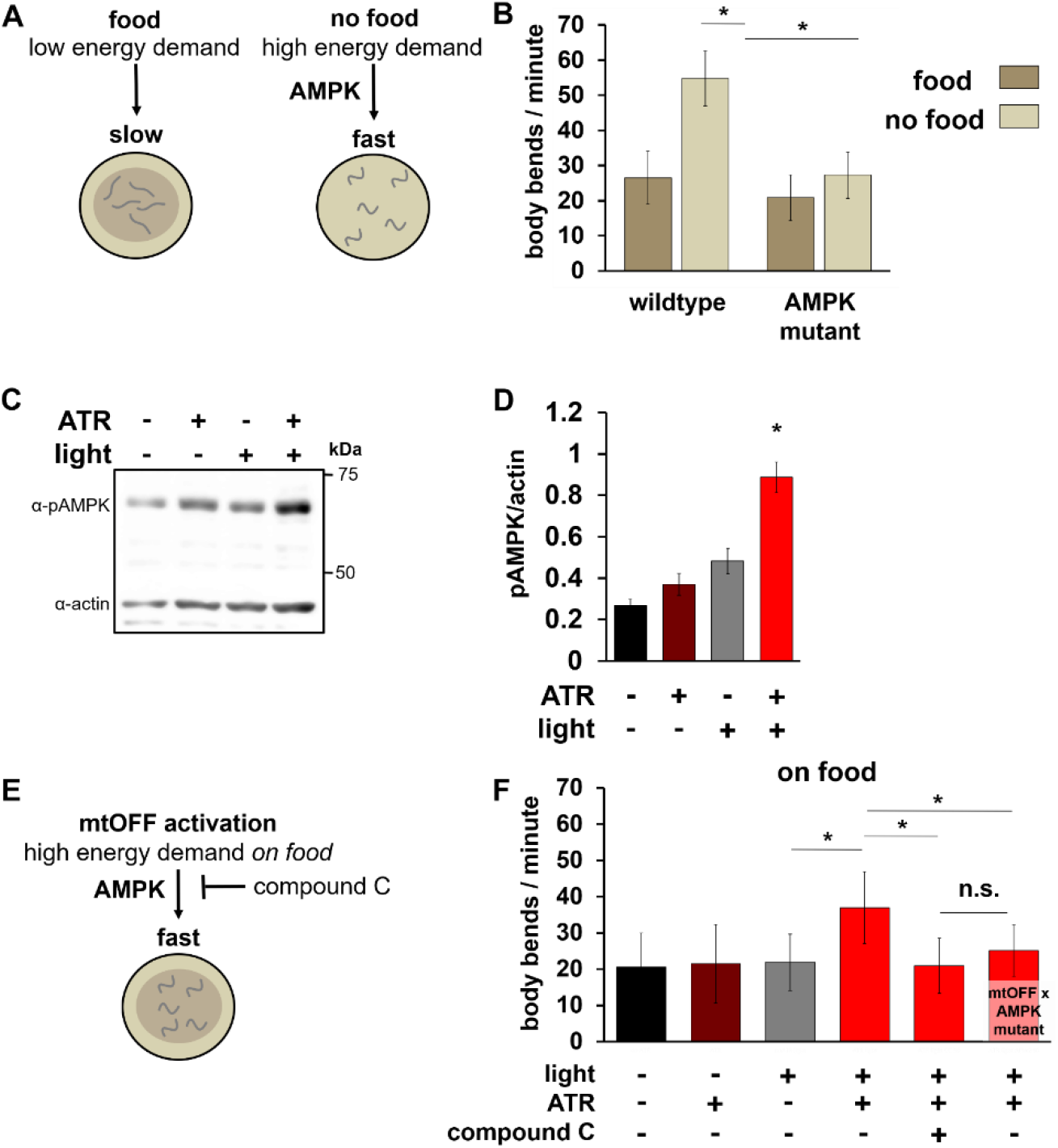
mtOFF modulates energy sensing behavior through AMPK. **A)** Schematic showing locomotion differences in *C. elegans* under both fed (left) and starved (right) conditions. Removal from food results in increased locomotion, mediated by AMPK signaling. This output is used in this study to validate functional AMPK signaling. **B)** Locomotion was scored by counting body bends per minute. Animals were scored either on food or immediately after being transferred off of food. *C. elegans* AMPK is encoded by the *aak-2* gene. The non-functional *aak-2(ok524)* mutant strain was used for AMPK mutant animals. One-way ANOVA with Tukey’s test for multiple comparisons was performed, wildtype on food vs wildtype off of food *p < 0.0001, wildtype off of food vs AMPK mutant off of food *p < 0.0001. Data are means ± SEM, n = 30 – 60 animals each condition from at least two experimental days. **C)** Immunoblot against phosphorylated (active) AMPK (pAMPK, top bands, 62 kDa) and actin (bottom bands, 43 kDa) from whole animal lysate on the same blot shows increased phosphorylation level under conditions of activated mtOFF. **D)** Quantification of pAMPK/actin densitometry shows increased pAMPK in response to mtOFF activation. pAMPK/actin is used to measure activated AMPK as there is no validated total AMPK antibody in *C. elegans*. One-way ANOVA with Tukey’s test for multiple comparisons was performed, -ATR -light vs +ATR +light *p = 0.0001, +ATR -light vs +ATR +light *p = 0.0006, -ATR +light vs +ATR +light *p = 0.0043. Data are means ± SEM, n = 4 independent blots from one plate of worms for each condition, with at least 1000 animals per plate. **E)** Schematic showing effects of mtOFF activation on locomotion. mtOFF is expected to create an energy demand through PMF dissipation that will increase locomotion, mediated by AMPK signaling. **F)** Body bends were scored, and illumination was throughout measurement where indicated. Animals were exposed to 50 μM compound C for 24 hours where indicated. One-way ANOVA with Tukey’s test for multiple comparisons was performed, -ATR +light vs +ATR +light *p < 0.0001, +ATR +light vs +ATR +light +compound C *p < 0.0001, +ATR +light vs mtOFF x AMPK mutant +ATR +light +compound C *p < 0.0001. n.s. is not significant, p = 0.3999. Data are means ± SEM, n = 30 – 60 animals each condition from at least two experimental days.

AMPK signaling in neurons alone is sufficient for driving increased locomotion in response to energy demand (Ahmadi and Roy, 2016). Therefore, we tested whether mtOFF could trigger increased locomotion with AMPK signaling only functional in neurons to signal perceived metabolic demand. Using a pan-neuronal gene promoter, we rescued neuronal AMPK expression in AMPK mutant animals as previously described (Ahmadi and Roy, 2016) and confirmed its sufficiency for restoring increased locomotion upon starvation. AMPK mutant animals expressing mtOFF did not respond to starvation by increasing locomotion to the same degree as wildtype, unless AMPK was rescued in neurons (Figure 3A). Under fed conditions, mtOFF activation increased locomotion in AMPK mutant animals with rescued neuronal AMPK (Figure 3B), similar to our results with mtOFF in a wildtype background. This confirmed that neuronal AMPK alone was sufficient to respond to dissipated PMF. Following this result in neurons, we asked whether tissue-restricted mtOFF activity could have similar effects on locomotion. We tested this hypothesis using tissue-specific gene promoters to drive mtOFF expression separately in neurons and in intestine (Figure 3C). We chose to also test intestinal expression of mtOFF to rule out the possibility of signaling coming from intestine, the organ that may sense energy availability through its nutrient-absorbing function, creating the internal metabolic state. Using a pan-neuronal gene promoter (*rab-3p*) and an intestinal gene promoter (*vha-6p*) to drive mtOFF expression, we found that neuronal mtOFF activation was sufficient to trigger increased locomotion (Figure 3D), but intestinal mtOFF activation was not (Figure 3E). In both cases, light exposure alone resulted in small increases in locomotion, however, under conditions of active mtOFF (light plus ATR) only neuronal mtOFF increased locomotion. These results indicate that mitochondrial function and energy sensing through AMPK are tightly linked and especially important in neurons. This tight spatial control of mitochondria and subsequent control of behavior shows that communication throughout the cell can be traced back to PMF changes in mitochondria.

**Figure 3.**
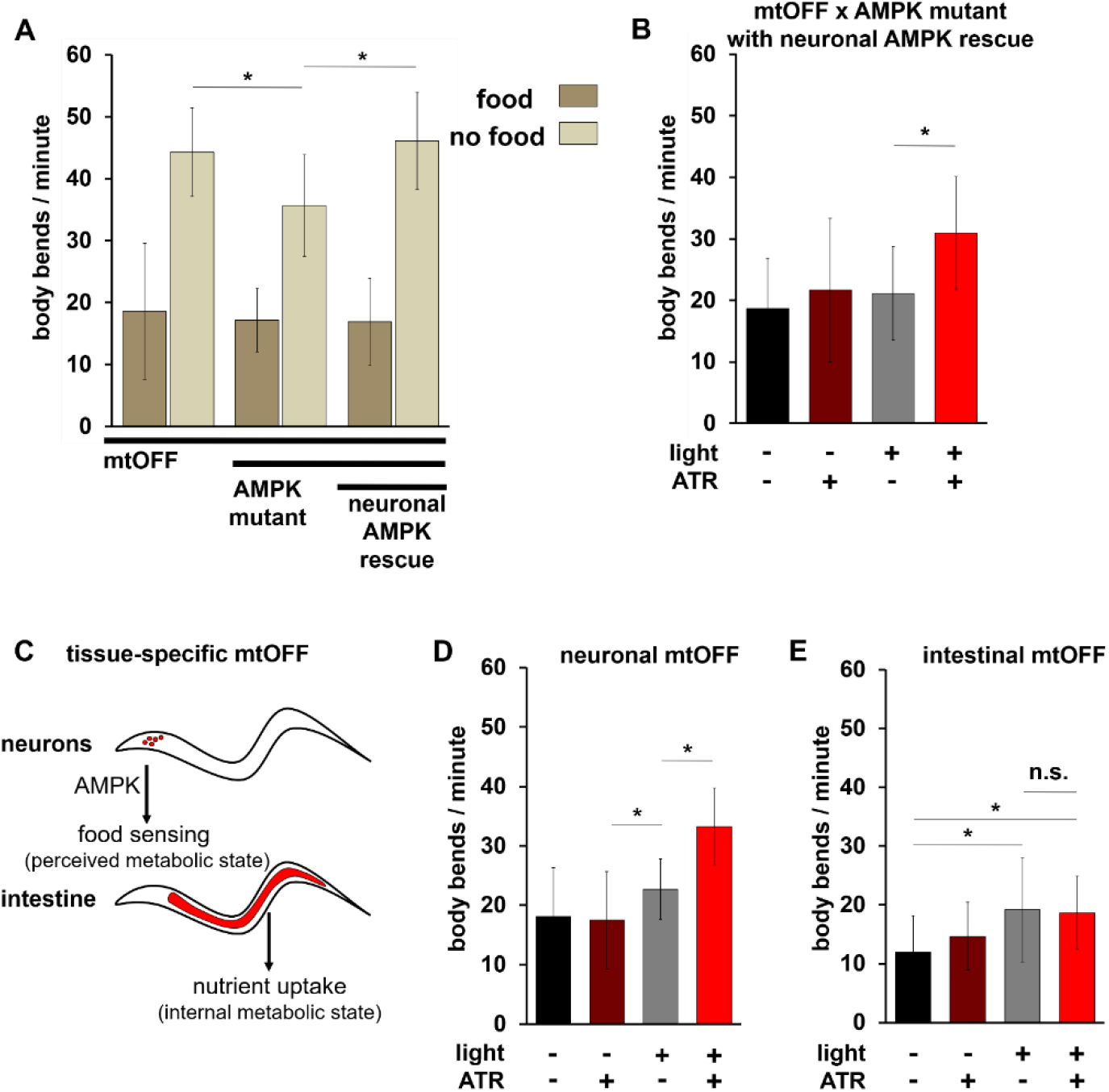
mtOFF triggers neuronal AMPK to control locomotion. **A)** Locomotion was scored by counting body bends per minute. Animals were scored either on food or immediately after being transferred off of food. All animals are expressing mtOFF, the center pair of bars are AMPK mutant animals, and the last pair of bars are AMPK mutant animals with functional AMPK re-expressed in neurons alone. AMPK mutant animals have a blunted response to starvation, and neuronal AMPK is sufficient to restore the response. One-way ANOVA with Tukey’s test for multiple comparisons was performed, mtOFF off of food vs mtOFF x AMPK mutant off of food *p < 0.0001, mtOFF x AMPK mutant off of food vs neuronal AMPK rescue *p < 0.0001. n.s. is not significant, p = 0.9910. Data are means ± SEM, n = 30 – 60 animals each condition from at least two experimental days. **B)** Locomotion in AMPK mutant animals expressing mtOFF with functional AMPK expressed only in neurons. Illumination was throughout measurement where indicated. mtOFF activation increased locomotion with functional AMPK expressed in neurons. One-way ANOVA with Tukey’s test for multiple comparisons, -ATR +light vs +ATR +light, *p < 0.0001. Data are means ± SEM, n = 30 – 60 animals each condition from at least two experimental days. **C)** Schematic showing tissue-specific mtOFF expression in neurons alone or in intestine alone. Neurons are responsible locomotion response to food sensation in an AMPK dependent manner. mtOFF was expressed in neurons to test if PMF loss in neurons alone could mediate an increased locomotion response. Intestine is the organ that absorbs nutrients, and mtOFF was expressed here to rule out the role of intestinal control of locomotion in response to PMF loss. **D)** Locomotion in animals expressing mtOFF only in neurons. Illumination was throughout body bends measurement where indicated. mtOFF activation in neurons increased locomotion compared to controls. Light alone also increased locomotion. One-way ANOVA with Tukey’s test for multiple comparisons was performed, -ATR vs -ATR +light *p = 0.0251, +ATR -light vs -ATR +light *p < 0.0083, -ATR +light vs +ATR +light *p < 0.0001. Data are means ± SEM, n = 30 – 60 animals each condition from at least two experimental days. **E)** Locomotion in animals expressing mtOFF only in intestine. Illumination was throughout body bends measurement where indicated. mtOFF activation in intestine did not increase locomotion compared to controls. Light alone also increased locomotion, similar to panel D. One-way ANOVA with Tukey’s test for multiple comparisons was performed, -ATR -light vs -ATR +light *p = 0.0007, -ATR -light vs +ATR +light *p = 0.0019. n.s. is not significant, p = 0.9910. Data are means ± SEM, n = 30 – 60 animals each condition from at least two experimental days.

### mtOFF protects against hypoxia through neuronal AMPK

Based on the link between energy availability and stress resistance, we next tested how changes in mitochondrial function elicit stress resistance *in vivo*. An animal’s perceived or internal energetic state can influence responses to hypoxia through many mechanisms, one example being AMPK signaling in fed versus starved nutritional states (LaRue and Padilla, 2011). AMPK activity in neurons alone is a model of perceived metabolic state to elicit a fast response in whole *C. elegans* (Ahmadi and Roy, 2016). Low nutrition or starvation causes stress resistance and specifically hypoxia resistance (Iranon et al., 2019, Scott et al., 2002). It follows that AMPK is required in many models of hypoxia resistance (LaRue and Padilla, 2011, Nishino et al., 2004, Chen et al., 2018, Emerling et al., 2009) as AMPK is a sensor of metabolic state. Because our optogenetic approach allows for spatiotemporal control of mitochondrial function, we therefore tested the ability of prophylactic PMF dissipation to protect against hypoxia (Figure 4A) and asked if neuronal AMPK could be specifically involved. Chronic PMF dissipation through pharmacology and genetics protects against hypoxia (Otani, 2004, Ozcan et al., 2013, Hoerter et al., 2004, Sack, 2006, Korde et al., 2005), however, PMF dissipation may only be required before hypoxia exposure for some mechanisms of resistance (Berry et al., 2020b). Therefore, we tested if mtOFF activation could protect *C. elegans* against impending hypoxia. Consistent with mammalian cell models, mtOFF activation to dissipate the PMF was also protective in our *C. elegans* model (Figure 4B) when we measured the percent of increased survival after hypoxia and light treatment, either with or without ATR. In addition, AMPK was required for that protection, as mtOFF activation in AMPK mutant animals was not protective (Figure 4C). We then tested if neuron-specific AMPK rescue could restore protection and found that mtOFF activation was again protective (Figure 4D). These data indicated neuronal AMPK activity was sufficient for hypoxia resistance. This supports a model of perceived metabolic state through neuronal AMPK can act to signal organism-wide protection.

**Figure 4.**
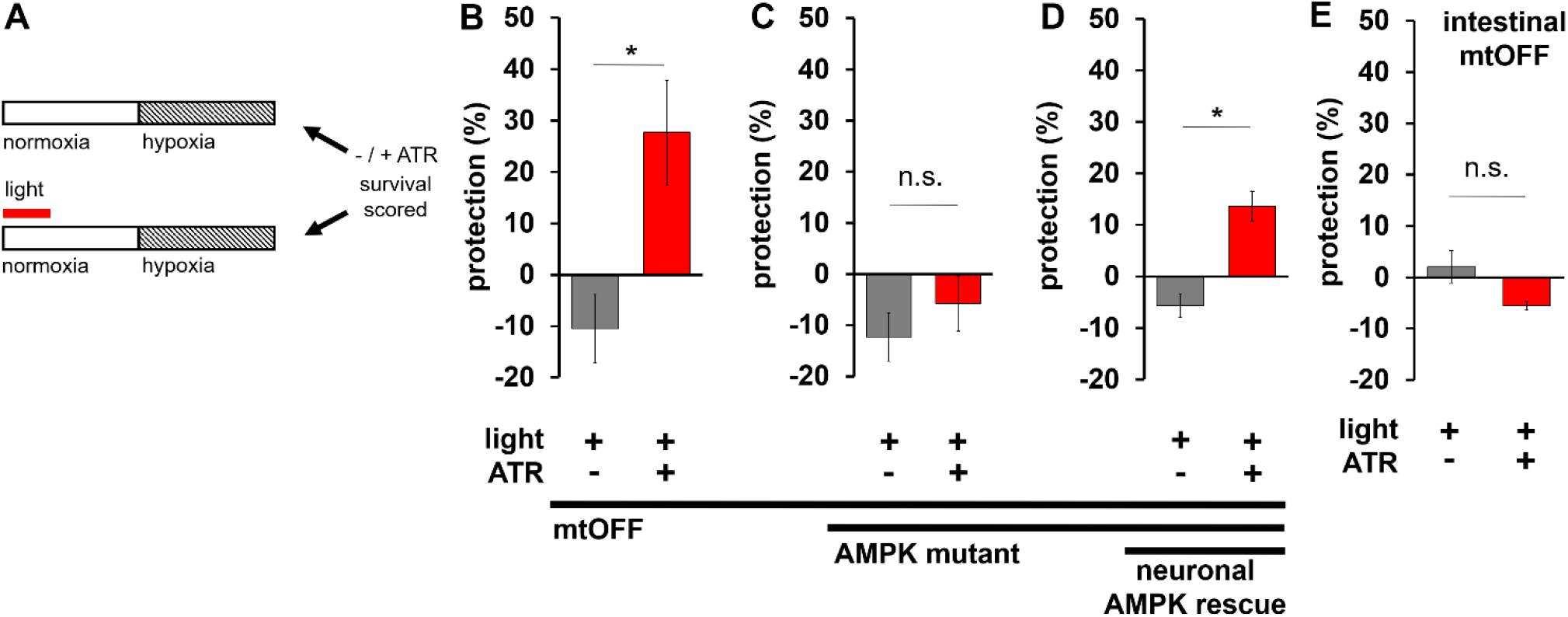
mtOFF protects against hypoxia through neuronal AMPK. **A)** Schematic showing protocol to activate mtOFF before hypoxia exposure to test the prophylactic effects of PMF dissipation on hypoxia resistance. Control conditions with and without ATR were either treated with light or left in the dark before hypoxia exposure. Normoxia is denoted with an open bar, and hypoxia is denoted by a striped bar. Timeline not to scale (see methods). To assess protection against hypoxia, the percent survival 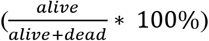 under -light conditions was subtracted from percent survival under +light conditions with and without ATR (top bar subtracted from bottom bar). The resulting protection percent (%) would be negative for damaging interventions after hypoxia exposure, and positive for interventions protective against hypoxia. Experiments are paired by concurrent hypoxia exposure. **B)** mtOFF activation prior to hypoxia conferred protection. Two-tailed paired t test, *p = 0.0082. Data are means ± SEM, n = 5, where one n is an average of three technical replicates of plates containing 15-50 animals. **C)** AMPK mutant animals expressing mtOFF were not protected against hypoxia, suggesting the protection observed in panel B requires AMPK signaling. Two-tailed paired t test was performed, n.s. is not significant, p = 0.176. Data are means ± SEM, n = 4-5, where one n is an average of three technical replicates of plates containing 15-50 animals. **D)** AMPK mutant animals expressing mtOFF with functional AMPK expressed in neurons alone conferred protection against hypoxia, suggesting AMPK activity in neurons alone is sufficient for the hypoxia resistance triggered by decreased PMF. Two-tailed paired t test was performed, *p = 0.0069. Data are means ± SEM, n = 3, where one n is an average of three technical replicates of plates containing 15-50 animals. **E)** Intestinal mtOFF activation prior to hypoxia conferred protection. Two-tailed paired t test was performed, n.s. is not significant, p = 0.645. Data are means ± SEM, n = 3, where one n is an average of three technical replicates of plates containing 15-50 animals.

In lifespan studies, and in the context of mitochondrial proteostatic stress, intestinal mitochondria mediate robust stress resistance (Durieux et al., 2011) which can act in the absence of neuronal mitochondrial dysfunction (Zhang et al., 2018). We therefore sought to test if direct intestinal PMF dissipation could result in resistance against hypoxia. In animals expressing mtOFF only in intestine, mtOFF activation did not confer protection against hypoxia (Figure 4E). These data show acute PMF loss in intestine specifically is not sufficient for hypoxia resistance. While proteostatic maintenance in mitochondria mediates stress resistance within intestine (Durieux et al., 2011, Zhang et al., 2018, Pena et al., 2016), direct PMF changes do not play a role in this tissue. This result suggests that in the absence of perceived energy crisis through neuronal AMPK activity, there is also absence of organism-wide stress resistance. In summary, our data suggest that PMF dissipation is a trigger for AMPK-mediated hypoxia resistance, and that neuronal AMPK is a driver of whole organism protection.

## DISCUSSION

In line with what is known about trans-tissue signaling in mitochondrial stress resistance (Durieux et al., 2011, Zhang et al., 2018, Pena et al., 2016), neuronal activation of AMPK conferred stress resistance (Figure 4B-D). We found that direct PMF loss in the intestine is not sufficient for protection (Figure 4E). In other words, intestine-triggered protection requires an external signal; acute PMF loss in intestine is not sufficient. This is consistent with studies on proteostatic mechanisms of hypoxia resistance (Durieux et al., 2011, Zhang et al., 2018, Pena et al., 2016, Kaufman and Crowder, 2015). Activation of mitochondrial unfolded protein responses in intestine drives protection, not necessarily acute mitochondrial PMF loss in intestine. Hypoxia-protective unfolded protein responses in intestine requires specific neuronal signals (Pena et al., 2016, Zhang et al., 2018), and here we show that neuronal AMPK signaling is also sufficient for organismal stress resistance (Figure 4B-D). We also show that the neuronal AMPK signaling can be activated directly by PMF dissipation in living organisms.

AMPK activity controls hypoxia resistance differently depending on nutritional quality (LaRue and Padilla, 2011), linking energy sensing with hypoxia resistance. AMPK activity in neurons is sufficient for behavioral responses to energy availability as well (Figure 3A) (Ahmadi and Roy, 2016); we add to these findings by showing neuronal AMPK control of behavior can be achieved through PMF dissipation in neurons (Figure 3D). AMPK activity in neurons alone models animal perception of metabolic state, and this signal alone can alter physiologic responses (Ahmadi and Roy, 2016). Our findings therefore suggest that mitochondrial energetics in neurons can result in these organismal responses as well, through AMPK. This again highlights the importance of neuronal signaling throughout organisms to respond to environmental stress.

It is well-established that changes in mitochondrial function can signal throughout organisms to extend *C. elegans* lifespan (Durieux et al., 2011). In addition, neuronal AMPK mediates longevity in *C. elegans*, and is interconnected with other metabolic pathways that affect mitochondrial morphology (Zhang et al., 2019). Our results complement each of these findings and suggest that the PMF may be the source change that leads to signaling downstream to affect physiology. We have shown that cellular energy sensing and responses to oxygen availability are both influenced by changes in the PMF. Together, our study and the data of others combine to say that fundamental changes in mitochondrial PMF trigger cellular signaling that can act throughout organisms to prepare for impending stress. This coordinated signaling likely occurs in a network of parallel and related pathways that fundamentally regulate and fine tune metabolism to adapt to environmental changes. Our approach to link energy sensing, hypoxia resistance and mitochondrial function was precise in space and time due to the application of optogenetics. Restricting light activation of mtOFF to before hypoxia alone, conferred temporal control that allowed testing PMF dissipation as a preventive intervention. Expressing mtOFF tissue specifically conferred spatial control over mitochondrial function. We were able to isolate the PMF as an independent variable in a living organism to link neuronal AMPK signaling to both behavior changes and to hypoxia resistance. We show these known signaling responses are linked through the PMF. This new link between phenomena demonstrates how mitochondria can integrate different aspects of cellular metabolism to signal adaptively, similar to our previous findings with mtON (Berry et al., 2020b). Changes in PMF have been correlated to different metabolic states and phenotypes, and here we move from correlation to causation through specific PMF control.

Using optogenetics to modulate PMF specifically has been recently applied in cellular models of disease to implicate other signaling pathways that are under mitochondrial PMF control, such as mitochondrial autophagy (Ernst et al., 2019), glucose stimulated ATP production in beta cells (Tkatch et al., 2017), muscle contraction through calcium signaling (Tkatch et al., 2017), and pathways of neurodegeneration (Imai et al., 2019). These results show that cellular activities under diverse metabolic control can each be manipulated by upstream intervention at one point, the PMF. Our development of mtOFF applied this optogenetic control in animals to begin to understand the signaling abilities of mitochondria *in vivo*. Fundamentally, these studies suggest that targeting mitochondria directly could be a more powerful intervention than targeting single proteins or pathways therapeutically; targeting mitochondria directly may modulate many signaling pathways concurrently for more powerful physiologic results. Our data also suggest that simply altering perceived metabolic state (through neuronal AMPK) should be characterized in other stress-resistance models. Overall, we show how mitochondrial PMF coordinates the stimuli of perceived energy availability and O_2_ levels, fundamental requirements for life, to prepare organisms to respond to stress.

## METHODS

### RESOURCE AVAILABILITY

#### Lead Contact

Further information and requests for resources and reagents should be directed to and will be fulfilled by the Lead Contact, Andrew Wojtovich.

#### Materials Availability

Materials generated in this study will be made available upon request to the Lead Contact, Andrew Wojtovich.

#### Data and Code Availability

This study did not generate/analyze datasets or code. All experimental data is presented in the manuscript, and raw data files will be made available upon request to the Lead Contact, Andrew Wojtovich.

### EXPERIMENTAL MODEL AND SUBJECT DETAILS

Adult hermaphrodite *Caenorhabditis elegans* were used in all experiments. Each experimental strain was backcrossed to wildtype at least 4 times. Animals were maintained at 20°C on nematode growth medium (NGM) seeded with OP50 *E. coli*.

### METHOD DETAILS

#### Molecular biology

*In vivo* transgene construction was carried out by homology-directed single-copy CRISPR/Cas9 gene insertion using the Mos1 mediated CRISPR Insertion (mmCRISPi) method (Philip et al., 2019). Briefly, transgenes were built through recombineering of 4 PCR fragments each containing at least 35 base pairs of homology. The fragments encoded a promoter, a mitochondrial targeting sequence of SDHC1 fused to the light activated proton pump, Mac, the red fluorescent protein mKate, and a characterized 5’ untranslated region from *unc-54*. The DNA coding sequence for 144 N terminal amino acids of the rat SDHC1 protein were fused to the N terminus of Mac by molecular cloning (Chow et al., 2010, Berry et al., 2020b) to achieve mitochondrial expression. The *eft-3* promoter was amplified from plasmid DNA pDD162 (forward amplification primer: ACAGCTAGCGCACCTTTGGTCTTTTA, reverse amplification primer: ACAACCGGTGAGCAAAGTGTTTCCCA). The *rab-3* promoter was amplified from plasmid DNA pSEP45 (Pena et al., 2016)(forward amplification primer: TCAGTGCAGTCAACATGTCGAGTTTCGTGCCGAATGACGACGACGACCTCGACGGCAAC, reverse amplification primer: GCCATTTTTAAGCCTGCTTTTTTGTACAAACTTGTCTGAAAATAGGGCTACTGTAG). The *vha-6* promoter was amplified from plasmid DNA pELA10 (Allman et al., 2009) (forward amplification primer: TCAGTGCAGTCAACATGTCGAGTTTCGTGCCGAATAGCACAGAACTGCATTAAG, reverse amplification primer: GCCATTTTTAAGCCTGCTTTTTTGTACAAACTTGTATTTTTATGGGTTTTGGTAG). SDHC1∷Mac was amplified from plasmid DNA pBB38 (forward amplification primer: ACAAGTTTGTACAAAAAAGCAGGCTTAAAAATGGCTGCGTTCTTGCTGAGAC, reverse amplification primer: GGATCCTCCTCCTCCAGATCCTCCTCCACCTCGGGCGCCGTCGTCCTCGCCGATC). mKate was amplified from plasmid DNA pAP088 (forward amplification primer: cccgaGGTGGAGGAGGATCTGGAGGAGGAGGATCCATGGTTTCCGAGTTGATCAAGG, reverse amplification primer: TTAACGATGTCCGAGCTTGGATGGGAGATCACAATATC). These PCR fragments amplified with unique homology regions, the first and fourth with homology to the genomic cut site, were microinjected into *C. elegans* hermaphrodite gonads with purified Cas9 protein and crRNA (GTCCGCGTTTGCTCTTTATT, DNA target) to achieve transgene construction into the genome of progeny such that gene promoters were followed by SDHC1∷Mac∷mKate∷3’ UTR when integrated in the genome. Transgenic lines generated by mmCRISPi were outcrossed to the wild‐type strain at least four times.

#### C. elegans *strains growth and maintenance*

Animals were maintained at 20°C on nematode growth medium (NGM) seeded with OP50 *E. coli*. All-*trans* retinal (ATR) was added to OP50 used for seeding NGM plates for a final concentration of 100 μM where indicated, as previously described (Berry et al., 2020b). Day 1 adult hermaphrodite animals were used for all experiments. Transgenic strains were generated by plasmid DNA microinjection as described (Mello and Fire, 1995), and the mmCRISPi method (Philip et al., 2019) as described above. For a complete strain list, see Supplementary Table 1.

#### Fluorescence microscopy

Images were obtained using a FV1000 Olympus laser scanning confocal microscope using a 60× oil objective (Olympus, N.A. 1.42), 561 nm diode laser illumination for red fluorescence and 488 nm for green fluorescence. Where indicated, animals were stained with 12 μM MitoTracker Green FM (Thermo Fisher Scientific, Waltham, MA) diluted in DMSO into the OP50 food for 20 hours (DMSO < 0.02% final). Line scan pixel intensity was performed using ImageJ software. Fluorescent determination of mtOFF localization was performed as previously reported (Trewin et al., 2019, Berry et al., 2020b, John et al., 2005). Briefly, cross‐section intensity plots of mitochondrial fluorescence (coexpressing either mtOFF∷mKate and GFP, or mtOFF animals stained with MitoTracker) were smoothed by three‐point moving averages and then normalized to maximum intensity. Distance between inflection points (defined as a threshold of 10% increase in pixel intensity from the previous point, in the direction from outer border toward the middle of the mitochondrion) was measured in pixels and converted to μm.

#### Mitochondria isolation

*C. elegans* mitochondria were isolated from day 1 adult animals using differential centrifugation as previously described (Wojtovich et al., 2011, Berry et al., 2020b). Briefly, fed animals from 3 to 5 15‐cm culture plates (~1 million animals) were transferred into 50 mL of M9 media (22 mM KH_2_PO_4_, 42 mM Na_2_HPO_4_, 86 mM NaCl, 1 mM MgSO_4_, pH 7) in a conical tube and settled by gravity on ice. Animals were rinsed with ice‐cold M9 twice, then once with ice‐cold mitochondrial isolation media (220 mM mannitol, 70 mM sucrose, 5 mM MOPS, 2 mM EGTA, pH 7.4) with 0.04% BSA. After again settling by gravity, the supernatant was removed, and worms were transferred onto ~2 g of pure sea sand per 1 mL of animals in an ice‐cold mortar. Animals were ground with an ice‐cold pestle for 1 min and extracted from the sand using mitochondrial isolation media and transferred to a 10‐mL conical tube. The samples were then homogenized in an ice‐cold glass Dounce homogenizer with 40 strokes. The homogenate was centrifuged at 600 g for 5 min, then the supernatant was transferred to a new tube and centrifuged at 700 g for 10 min. The second pellet was resuspended in 1 mL of mitochondrial isolation media without BSA in a 1.5 mL tube and centrifuged at 7,000 g for 5 min. The final pellet was resuspended in 50 μL of mitochondrial isolation media without BSA. Protein was quantified using the Folin‐phenol method.

#### Light sources for mtOFF activation

Illumination sources were a 580 nm Quantum SpectraLife LED Hybrid lamp by Quantum Devices, Barneveld WI (abbreviated Quantum LED in the text), a 540-600 nm GYX module, X-Cite LED1 by Excelitas, Waltham MA, (abbreviated XCite LED), and a 540-580 nm excitation filter MVX10 Fluorescence MacroZoom dissecting microscope by Olympus (abbreviated MVX) powered by an X-Cite 220 V mercury bulb by Excelitas. Light intensities are indicated for each experimental condition and were determined with a calibrated thermopile detector (818P-010-12, Newport Corporation, Irvine, CA) and optical power meter (1916-R, Newport Corporation).

#### Immunoblotting

Fed adult animals were harvested in M9 media after 4 hour exposure to 1 Hz light (Quantum LED, 0.02 mW/mm^2^). Animals were centrifuged at 1,000 g for 1 min and ground by plastic pestle disruption in lysis buffer (20 mM Tris–HCl, 100 mM NaCl, 1 mM EDTA, 1 mM DTT, 10% glycerol, 0.1% SDS, pH 7.6, 1× Halt protease inhibitor cocktail, Thermo78429). Samples were then diluted 1:1 in sample loading buffer (100 mM Tris–HCl, 10% glycerol, 10% SDS, 0.2% w/v bromophenol blue, 2% β‐mercaptoethanol). Samples were heated at 95°C for 5 min, and 12.5 μg of protein was loaded in each lane of a 7.5% polyacrylamide gel for separation by SDS–PAGE. Protein was transferred to nitrocellulose membranes, blocked using 5% non‐fat milk/TBST (50 mM Tris, 150 mM NaCl, 0.05% Tween 20, pH 8.0) for 1 h at room temperature, and incubated at 4°C in primary antibodies diluted 1:1,000 in 5% bovine serum albumin. Membranes were washed in TBST and incubated in horseradish peroxidase‐conjugated secondary antibodies for 1 h at room temperature. Antibodies used: 1:2000 anti‐Actin (Abcam #ab14128), 1:10,000 anti‐phospho‐AMPKα Rabbit (Cell Signaling, #2535), 1:2,000 anti‐rabbit IgG (Cell Signaling #7074S), and anti‐mouse IgG (Thermo Scientific #32430). Detected proteins were visualized by chemiluminescence (ChemiDoc, Bio‐Rad) using ECL (Clarity Western ECL Substrate, Bio‐Rad). Densitometry was performed using ImageJ software.

#### Mitochondrial membrane potential measurement

0.5 mg/mL isolated mitochondria were stirred in mitochondrial respiration buffer (MRB: 120 mM KCl, 25 mM sucrose, 5 mM MgCl_2_, 5 mM KH_2_PO_4_, 1 mM EGTA, 10 mM HEPES, 1 mg/ml FF‐BSA, pH 7.35) at 25°C with 2 μM rotenone and 5 mM succinate. 20 nM tetramethylrhodamine ethyl ester (TMRE, Thermo Fisher, T669) was added to observe mitochondrial membrane potential in non-quench mode, where TMRE accumulates in the matrix and fluorescence is high in the presence of mitochondrial membrane potential. Upon addition of the protonophore FCCP, TMRE exits mitochondria and fluorescence decreases. TMRE signal was measured by Cary Eclipse Fluorescence Spectrophotometer (Agilent Technologies) using a 335–620 nm excitation filter and a 550–1,100 nm emission. Illumination was continuous throughout all measurements (0.39 mW/mm^2^ XCite LED) with increasing light dose (fluence, J/cm^2^). 2 μM FCCP was added to completely depolarize mitochondria and observe minimum fluorescence. Data are normalized to maximum succinate-fueled fluorescence (F/F_max_).

#### Mitochondrial matrix pH measurement

BCECF‐AM (Thermo Fisher, B1170), a ratiometric pH indicator, was used to measure pH change in the mitochondrial matrix (Jung et al., 1989, Aldakkak et al., 2010, Berry et al., 2020b) in response to mtOFF activation. Isolated mitochondria (~200 μL) were incubated with 50 μM BCECF‐AM for 10 min at room temperature. Mitochondria were pelleted at 7,000 g for 5 min at 4°C, isolation media was replaced and mitochondria were pelleted again to remove residual BCECF‐AM. Mitochondria were then assayed as described in the mitochondrial membrane potential measurements. 440 and 490 nm excitation wavelengths were used to measure 545 nm emission fluorescence using a Cary Eclipse Fluorescence Spectrophotometer (Agilent Technologies). Fluorescence ratio at 545 nm of 490/440 nm excitation wavelengths is presented to show pH changes in the mitochondrial matrix. Light treatment was 0.39 mW/mm^2^ (XCite LED), and 2 μM FCCP was used at the end of each trace to establish minimum signal. Change in the ratio (ΔBCECF ratio) value is presented comparing before and after illumination of mtOFF.

#### C. elegans *O*_2_ consumption

Whole animal O_2_ consumption was recorded using a Clark-type O_2_ electrode (S1 electrode disc, DW2/2 electrode chamber and Oxy-Lab control unit, Hansatech Instruments, Norfolk UK). Adult animals were collected in M9 and pelleted by centrifugation. Animals were then rinsed in M9 and added to the chamber in 0.5 mL of continuously stirred M9. Continuous light exposure (XCite LED) was 0.39 mW/mm^2^ throughout baseline measurement where indicated. 160 μM final concentration FCCP was added to induce maximal respiration, and 40 mM final concentration sodium azide was added to inhibit mitochondrial respiration. O_2_ consumption rates (baseline, maximal, and inhibited) were measured for 10 minutes or until stable. Animals were collected after measurement for protein quantification by the Folin-phenol method.

#### ATP measurements

Whole animals were used for ATP quantification by luciferase bioluminescence. Adult animals on OP50 seeded plates were exposed to 4 hours of 1 Hz light (Quantum LED, 0.02 mW/mm^2^), and then collected in M9 media in 1.5 mL tubes. Samples were quickly centrifuged to pellet animals and supernantant was removed to leave pelleted animals in 100 μL M9. Samples were freeze-cracked three times in liquid nitrogen and protein concentration was measured using the Folin-phenol method. Samples were then boiled at 100°C for 15 minutes, then placed on ice for 5 minutes. Samples were centrifuged at 14.8 g for 10 min at 4°C. An ATP determination kit was used according to the manufacturer’s instructions to measure ATP levels. (Invitrogen Molecular Probes, A22066). Fold change of relative ATP levels are presented after normalization to either – or + ATR baseline levels without light.

#### Locomotion measurement

The number of animal body bends in 15 seconds was scored on and off of OP50 food to assess locomotion (Sawin et al., 2000) (n = 30–60 animals scored on at least 2 separate days). Change in direction of motion of the posterior pharyngeal bulb was counted as one body bend (Tsalik and Hobert, 2003). AMPK inhibition was achieved by 24-hour exposure to 50 μM final concentration of compound C in the NGM plates. Where indicated, light treatment was continuous throughout body bend measurements (MVX, 0.265 mW/mm^2^).

#### Hypoxia exposure

A hypoxic chamber (Coy Laboratory Products, 5%/95% H_2_/N_2_ gas, palladium catalyst) was used at 26°C with 15-50 animals per plate for hypoxia experiments. O_2_ concentration was monitored and always < 0.01%. 1 Hz light exposure (Quantum LED, 0.02 mW/mm^2^) was applied for 4 hours, 20 hours before hypoxia exposure based on a time window identified for protective signaling to occur (Berry et al., 2020b). 18.5-hour hypoxic exposure was used to kill at least 50% of animals. 24 hours after hypoxia exposure moving animals or animals that moved in response to a light touch with an eyelash were scored as alive. Animals supplemented with ATR laid eggs onto plates without ATR that were subsequently used as adults in hypoxia experiments to minimize potential effects of ATR supplementation (Supplementary Figure 4) (Berry et al., 2020b). Data are presented as protection (%), where baseline survival was subtracted from the survival of animals exposed to light to show potential damaging or protective effects as negative or positive values, respectively.

### QUANTIFICATION AND STATISTICAL ANALYSIS

#### Statistics

Using GraphPad Prism (v7), we performed two-tailed unpaired t-tests and one- or two-way ANOVA with post-hoc tests where appropriate. P < 0.05 was considered significantly different. For hypoxia experiments, we used two-tailed paired t-tests where samples were paired by hypoxia exposure. Throughout, n values are biological replicates: independent animal populations. Technical replicates are from one biological replicate exposed separately to experimental interventions. See figure legends for detailed statistical information.

## LIST OF ABBREVIATIONS

mtOFF: mitochondria-OFF
IM: inner mitochondrial membrane
PMF: protonmotive force
ETC: electron transport chain
AMPK: AMP-activated protein kinase
mtON: mitochondria-ON
mmCRISPi: Mos1 mediated CRISPR Insertion
NGM: nematode growth medium
ATR: all-*trans* retinal
ChR2: channelrhodopsin 2
IMS: intermembrane space
MTS: mitochondrial targeting sequence
TMRE: tetramethylrhodamine ethyl ester
ΔΨm: mitochondrial membrane potential
ΔpH: pH gradient

## ACKNOWLEDGEMENTS

A.P.W. is supported by grants from National Institutes of Health (R01 NS092558, R01 NS115906). B.J.B. is supported by an American Heart Association Predoctoral Fellowship (18PRE33990054). We thank the Mitochondrial Research & Innovation Group at University of Rochester Medical Center for helpful discussions and guidance. Some strains were provided by the CGC, which is funded by NIH Office of Research Infrastructure Programs (P40 OD010440).

## AUTHOR CONTRIBUTIONS

B.B. and A.P.W. designed the research, B.B. performed the research, analyzed the data and wrote the paper, A.B. performed the research.

## DECLEARATION OF INTERESTS

The authors declare there are no competing interests to disclose.

---

**Supplementary Figure 1.**
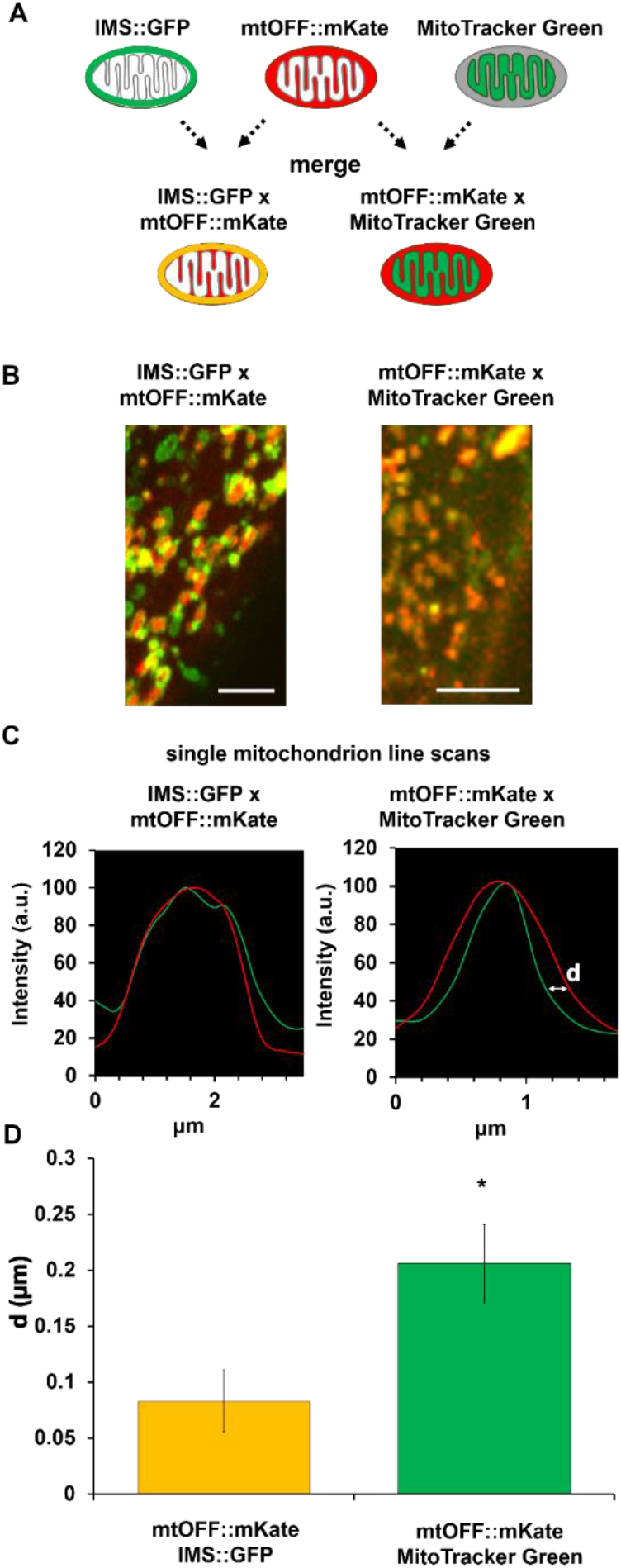
Mitochondrial localization of mtOFF. **A)** Schematic showing expected fluorescence localization in single mitochondria for mtOFF∷mKate, intermembrane space (IMS)∷GFP, and MitoTracker Green. Bottom row shows the expected fluorescence pattern of merged images. **B)** Fluorescent images of muscle mitochondria in live *C. elegans* coexpressing IMS∷GFP and mtOFF∷mKate (left) or expressing mtOFF∷mKate and stained with MitoTracker Green. Scale bars are 5 μm. **C)** Representative profile fluorescence intensity plots for single mitochondria from the images in panel B. The white letter d shows the distance between inflection points of the red and green fluorescent signals. **D)** The distance between inflection points was quantified (examples shown in panel d). mtOFF∷mKate localized close to IMS∷GFP signal, and distant from the matrix MitoTracker Green signal as expected, with the C terminal mKate predicted to be in the IMS. Two-tailed unpaired t test was performed, *p = 0.0137. Data are means ± SEM, n = 14 mitochondria from distinct animals for each condition.

**Supplementary Figure 2.**
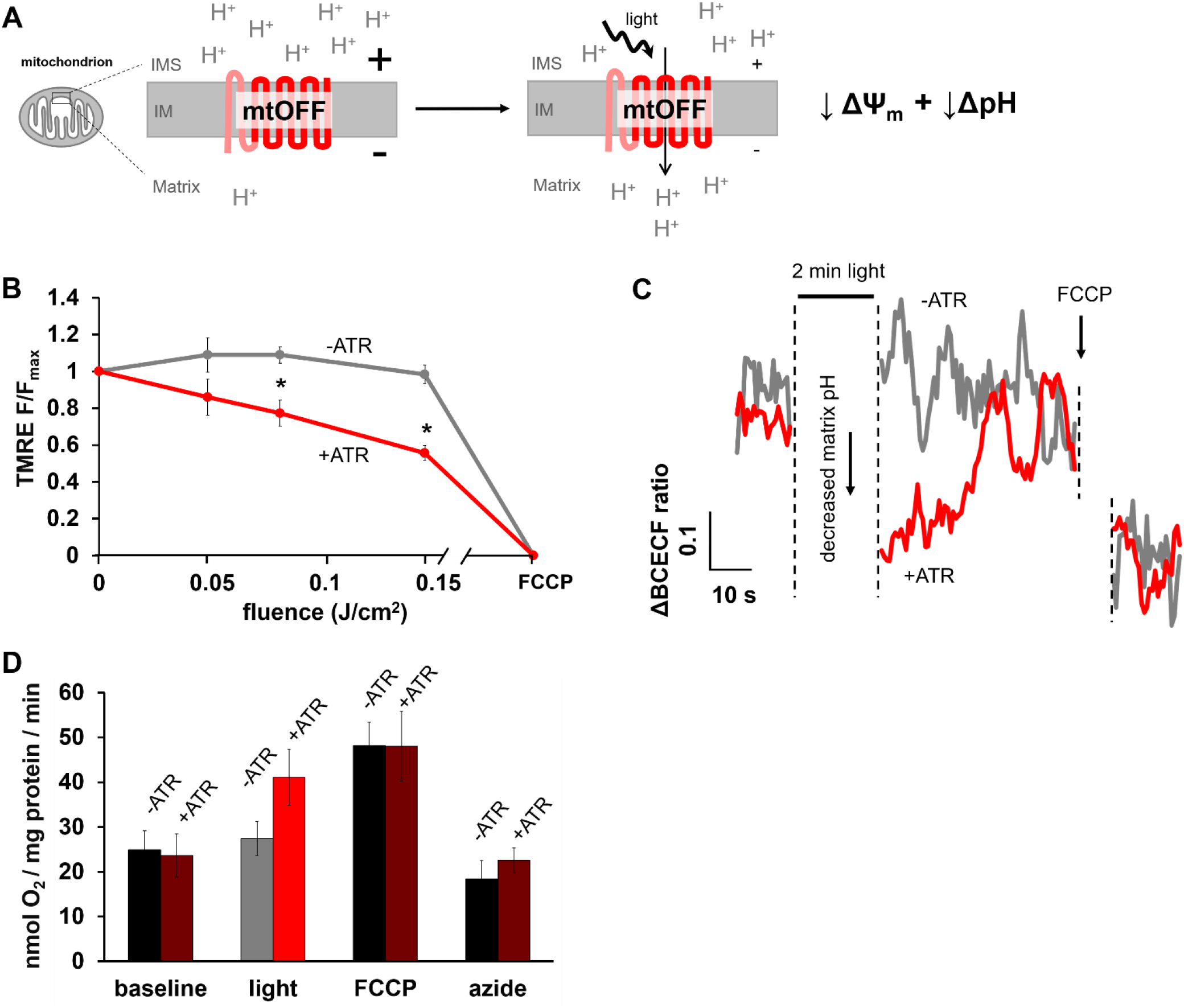
mtOFF decreases the PMF. **A)** Schematic showing mtOFF decreasing both components of the protonmotive force (PMF), the ΔΨ_m_ and the ΔpH, upon light exposure. mtOFF pumps protons (H+) from the intermembrane space (IMS) across the inner membrane (IM) into the matrix. **B)** TMRE fluorescence was measured in response to increasing light doses. Increasing fluence (light dose, Joules / cm^2^) results in progressively decreased PMF in isolated mitochondria supplied with succinate. Data from 0 and 0.15 J/cm^2^ are presented in Figure 1C. Two-way ANOVA with Holm-Sidak test for multiple comparisons was performed, 0.08 J/cm^2^ *p = 0.0068, 0.15 J/cm^2^ *p = 0.018, n = 4 independent mitochondrial isolations. **C)** Representative BCECF-AM fluorescence ratio trace. Baseline level of mitochondria supplied with succinate from animals with and without ATR is shown followed by light treatment (no BCECF-AM fluorescence measured), and signal immediately after illumination. Mitochondria with ATR have a decreased matrix pH, indicating proton entry through mtOFF during light exposure. Rapid reestablishment of baseline pH shows the reversibility of mtOFF when light is removed. FCCP was then added to establish minimum signal. **D)** Raw O_2_ consumption values under baseline, light treatment, maximal respiration, and minimum respiration states in whole animals. Maximum respiration was induced by FCCP treatment, and minimum respiration was induced with azide treatment. Data are presented for animals with and without ATR and are means ± SEM, n = 5, where one n is one O_2_ consumption rate from ~1500 animals in a Clark type O_2_ electrode. Normalized baseline data are presented in Figure 1E.

**Supplementary Figure 3.**
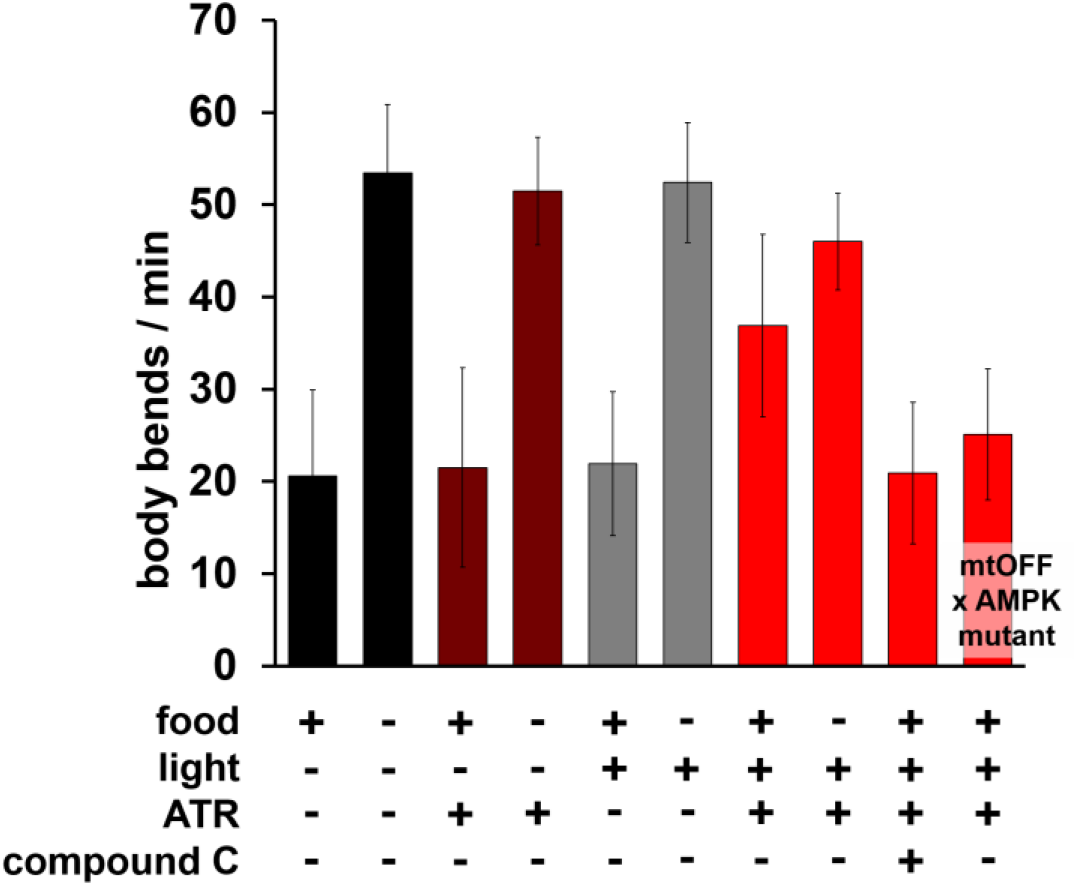
mtOFF does not affect starved locomotion speed. Graph includes locomotion rates from animals off of food, data from animals on food is presented here as well as in Figure 2E. Locomotion was not affected my mtOFF activation in animals off of food, as starvation creates an energy demand and increases locomotion, which is mimicked by mtOFF.

**Supplementary Figure 4.**
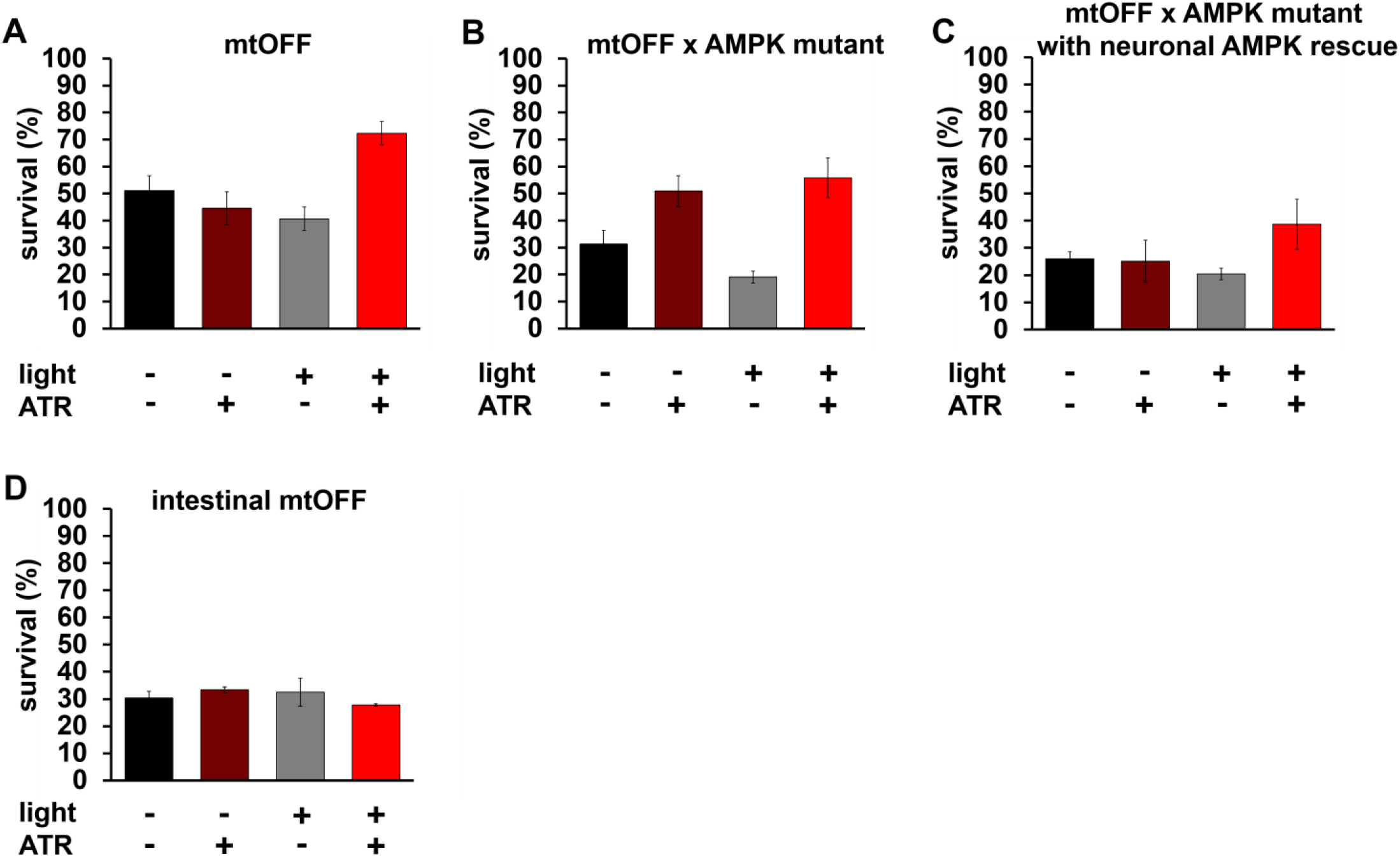
Raw survival data following hypoxia for Figure 4. These raw data were used to calculate protection % in Figure 4 to account for ATR and light exposure effects. See Figure 4 legend and methods section for details. **A)** Survival % for animals expressing mtOFF after hypoxia exposure. Protection % presented in Figure 4B. **B)** Survival % for animals expressing mtOFF in the AMPK mutant background after hypoxia exposure. Protection % presented in Figure 4C. **C)** Survival % for animals expressing mtOFF in the AMPK mutant background with AMPK expression rescued in neurons after hypoxia exposure. Protection % presented in Figure 4D. **D)** Survival % for animals expressing mtOFF in intestine after hypoxia exposure. Protection % presented in Figure 4E.

**Supplementary Table 1.** *C. elegans* strains.

| Strain | Genotype | Abbreviation | Source | Description |
| --- | --- | --- | --- | --- |
| N2(Bristol) |  | wildtype | *C. elegans* Genetics Center (CGC) | Wild type |
| APW208 | *jbmSi1[eft-3p::rat Sdhc1(144N')::Mac::mKate::unc-54 3'UTR \*ttTi4348] I* | mtOFF | this study | Ubiquitously expressed mitochondria-targeted light-activated proton pump, mtOFF. |
| RB754 | *aak-2(ok524) X.* | AMPK mutant | CGC | AAK-2 loss of function mutant from the *C. elegans* Gene Knockout Project at the Oklahoma Medical Research Foundation, which is part of the International *C. elegans* Gene Knockout Consortium. |
| APW240 | *jbmSi1[eft-3p::Sdhc1(144N')::Mac::mKate::unc-54 3'UTR \*ttTi4348] I; aak-2(ok524) X.* | mtOFF x AMPK mutant | this study | APW208 crossed to RB754 to achieve mtOFF expression in an AMPK loss-of-function background |
| APW254 | *jbmSi1[eft-3p::Sdhc1(144N')::Mac::mKate::unc-54 3'UTR \*ttTi4348] I; jbmEx105[pMR729(unc-119p::aak-2) 50 ng/uL, Pceh-22::GFP 2.5 ng/uL, Hyperladder 95 ng/uL]; aak-2(ok524) X.* | mtOFF x AMPK mutant with neuronal AMPK rescue | this study | APW240 expressing pan-neuronal AMPK |
| APW40 | *immt-1(jbm7 [immt-1::link::GFP]) X* | NA | (Trewin et al., 2019) | Strain expressing mitochondrial intermembrane space GFP used to generate APW252 |
| APW252 | *jbmSi1[eft-3p::Sdhc1(144N')::Mac::mKate::unc-54 3'UTR \*ttTi4348] I; immt-1(jbm7 [immt-1::link::GFP]) X* | IMS::GFP x mtOFF::mKate | this study | APW208 crossed with APW40 used for mtOFF localization and topology studies. |
| APW242 | *jbmSi5[vha-6p::Sdhc1(144N')::Mac::mKate::unc-54 3'UTR \*ttTi4348] I* | intestinal mtOFF | this study | Strain expressing intestinal mtOFF. |
| APW258 | *jbmSi9[rab-3p::Sdhc1(144N')::Mac::mKate::unc-54 3'UTR \*ttTi4348] I* | neuronal mtOFF | this study | Strain expressing pan-neuronal mtOFF. |

